# UDCT: Unsupervised data to content transformation with histogram-matching cycle-consistent generative adversarial networks

**DOI:** 10.1101/563734

**Authors:** Stephan Ihle, Andreas M. Reichmuth, Sophie Girardin, Hana Han, Flurin Stauffer, Anne Bonnin, Marco Stampanoni, János Vörös, Csaba Forró

## Abstract

The segmentation of images is a common task in a broad range of research fields. To tackle increasingly complex images, artificial intelligence (AI) based approaches have emerged to overcome the shortcomings of traditional feature detection methods. Owing to the fact that most AI research is made publicly accessible and programming the required algorithms is now possible in many popular languages, the use of such approaches is becoming widespread. However, these methods often require data labeled by the researcher to provide a training target for the algorithms to converge to the desired result. This labeling is a limiting factor in many cases and can become prohibitively time consuming. Inspired by Cycle-consistent Generative Adversarial Networks’ (cycleGAN) ability to perform style transfer, we outline a method whereby a computer generated set of images is used to segment the true images. We benchmark our unsupervised approach against a state-of-the-art supervised cell-counting network on the VGG Cells dataset and show that it is not only competitive but can also precisely locate individual cells. We demonstrate the power of this method by segmenting bright-field images of cell cultures, a live-dead assay of C.Elegans and X-ray-computed tomography of metallic nanowire meshes.

## Introduction

As modern imaging techniques are evolving at an ever faster pace, providing unprecedented automation capabilities, high resolution and ultra fast acquisition, scientists are often gathering more data than they can analyze. Be it wide area confocal laser microscopy time lapses, X-ray-computed tomographies, FIB- or Cryo-SEM of biological tissue, ultra-fast video acquisition of neural activity, there is a need for flexible image processing tools capable of handling large and complex datasets. For the task of image segmentation which consists in labeling regions of interest with a contour, color or intensity, deep convolutional neural networks[1] have proven to be extremely powerful and versatile. These algorithms are commonly used in a variety of different network architectures [2, 3, 4, 5] and their widespread use has been made possible in part by the open-source nature of AI research with codes being hosted in public repositories on GitHub, and also by user-friendly libraries such as PyTorch, Tensorflow, Keras and Caffe which allow to leverage the power of graphical processing units (GPUs) from a Matlab or Python environment. These networks can be thought of as intricate non-linear filters designed for a specific task which can, when applied to an image, highlight for example a tumor in a computerized tomography scan of lung tissue [6]. A standard way to design these task-specific networks is to provide a number of ground truth images (supervised learning) where a trained human labels the regions of interest. This practice is problematic as it requires a lot of human hours and it is not always clear how much data from the original set needs to be processed by the researcher to reliably train the computer. Lastly, in cases that require instance segmentation, for example fiber bundles where each individual fiber needs to be identified as opposed to just labeling the background and fibers distinctly [7], the human processing time becomes prohibitively expensive since overlapping fibers need to be labeled differently.

Here we propose a method that does not require human labeling of data (unsupervised learning).

We build on and enhance a previously published architecture named cycle-consistent generative adversarial networks (cycleGAN). The seminal publication by Zhu et. al [8] demonstrates how the algorithm can perform style transfer between two sets of images, turning winter landscapes into summery ones, Cezanne paintings into real photographs, and horses into zebras among many other examples. The architecture consists of two generative adversarial networks [9] (GAN) that are each trained to transfer an image from one domain into another. A GAN consists of a generator network and a discriminator network. The generator network is given an input image and produces another one using a convolutional neural network architecture. The discriminator is a classifier which receives an image and predicts whether it is genuine or created by a generator. In the cycleGAN, each generator takes images from its respective domain (horse or zebra) and creates images of the opposite (zebra or horse). While each discriminator is trained to distinguish generated images from real ones (generated horse or real horse), the generators in turn are trained to fool the discriminators. This adversarial procedure culminates in the creation of realistic images [10]. In order to ensure true style transfer, a cyclic constraint is enforced whereby the generated images are passed into the generators of the corresponding domain (a zebra generator is given a generated horse) and the result must be identical to the original image used to create the generated one (see Fig.1).

**Figure 1:**
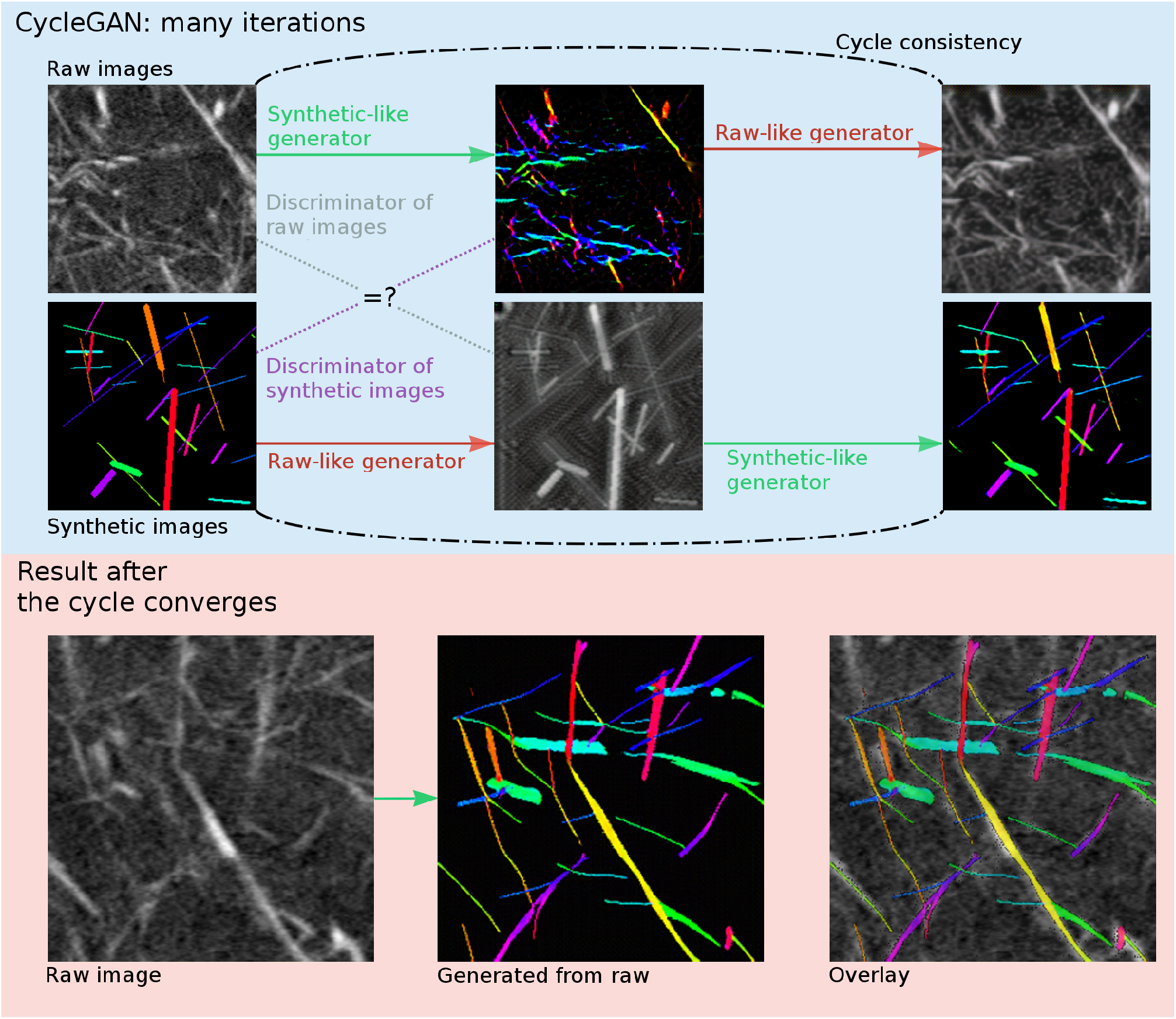
General UDCT procedure: we design computer generated data whose annotation encodes the information we wish to extract from the raw images, which in this case is individual wires. Two generator networks create images from the opposite image domain, with the goal to fool the corresponding discriminators. The networks are trained in a sequential fashion: the generators learn to generate images that can pass for genuine. Then, the discriminators are trained to tell apart the original and newly generated images. This procedure is repeated ideally until the generators create images that are so similar to the originals that the discriminators cannot, even with training, distinguish them. Additionally, during every generator training sequence, a cycle consistency is enforced, by penalizing discrepancies between a genuine image and its sequential application of the two generators. This ensures that the structure of the original images are preserved and only the style is modified. Once the network converges, the raw images are transformed by the generator that produces the synthetic-looking forgery and thereby a segmentation without ever requiring human data annotation.

An important aspect of the approach is that it does not require paired images and therefore in order to turn winter scenes into summer, one does not need a photograph of the same landscape in both seasons, but simply many summer and winter images. In the biomedical field we seek to turn real data into segmented data. Not needing paired images makes this approach very versatile, because one can use publicly available labeled data that share similarities with the images to segment [11, 12, 13, 14, 15]. However this still requires previously existing annotations and thus is a limiting factor. To overcome this, some attempts have been made to use synthetic data [16, 17] for cell-segmentation. We show that this approach has a much broader range of application, and use computer generated data that encodes the features we seek to highlight and by means of style transfer make the network turn the real images into segmented ones. We call our method unsupervised data to content transformation (UDCT).

We demonstrate the power of UDCT by evaluating it first on the common benchmark of counting cells in the public VGG dataset[18] and show that we are not only competitive with a state-of-the-art supervised network, we can extract, for example, the location of individual cells. We then move on to segment primary rat cortical neurons cultured in-vitro on glass slides. In this highly complex dataset containing large variations in cell-density and inconsistent image focus, we manage to distinguish alive from dead cells and outperform the same state-of-the-art supervised network. The next set consists of live-dead assay images of C.Elegans where we also investigate the importance of structured image backgrounds and their influence on convergence speed. Finally, we show a segmentation of an X-ray-computed tomography of silver nanowires embedded in an elastomer, which is a type of dataset that would be prohibitively expensive to label by hand if one wanted to achieve instance segmentation. Our own implementation of the cycleGAN and the data generating scripts and the evaluation of the results are all available at https://github.com/UDCTGAN/UDCT.

## Results

### VGG cell dataset

We demonstrate the validity of our approach by performing a counting task on a publicly available and commonly used dataset. We selected the VGG Cells [18] (Fig.2 raw images) because they are used for benchmarking deep convolutional approaches for counting cells and the state-of-the-art network, Count-ception, [19] is recent. Theirs is a supervised approach where the network learns to count objects knowing the true count of every image used during training. In order to show the versatility of our approach, we propose two ways of counting the cells without resorting to the ground truth labels. In a first synthetic dataset depicted in Fig.2a) we place circular objects, representing the cells, color-coded such that their overlaps are colored differently depending on whether they are the intersection of two, three or four cells. In a post-processing step, the colors are converted to values of 1 for cells, 2 for a pairwise intersection and 3 or 4 for three and fourfold overlaps. The pixels are then summed and divided by the average synthetic cell area to produce a count. This is similar to Cohen *et al*’s [19] state-of-the-art approach, with the distinction that ours is unsupervised. In a second dataset depicted in Fig.2b) we simply create round-shaped cells of varying intensity in the blue channel, a two-dimensional gaussian map with a standard deviation of 3 in the green and 1 in the red channel. A gaussian with a larger width is more easily mapped onto the cells of the raw images, since they look similar. However, the cells are better localized with a sharper gaussian. To simplify the task of mapping sharp gaussians to the images, we added both a sharp and wide gaussian to the synthetic dataset. Because of the similarity in shapes in both datasets, the network finds the mapping between synthetic and raw cells quickly, and the gaussian maps localize the center point of the cell. The latter are then reliably located in the transformed images by maximum likelihood estimation. Therefore, in addition to counting the cells, we can also retrieve their position in an unsupervised fashion. In both synthetic datasets, we randomly placed between 70 and 300 cells in each image. In Table 1 we summarize our results and compare them to the Count-ception network.

**Figure 2:**
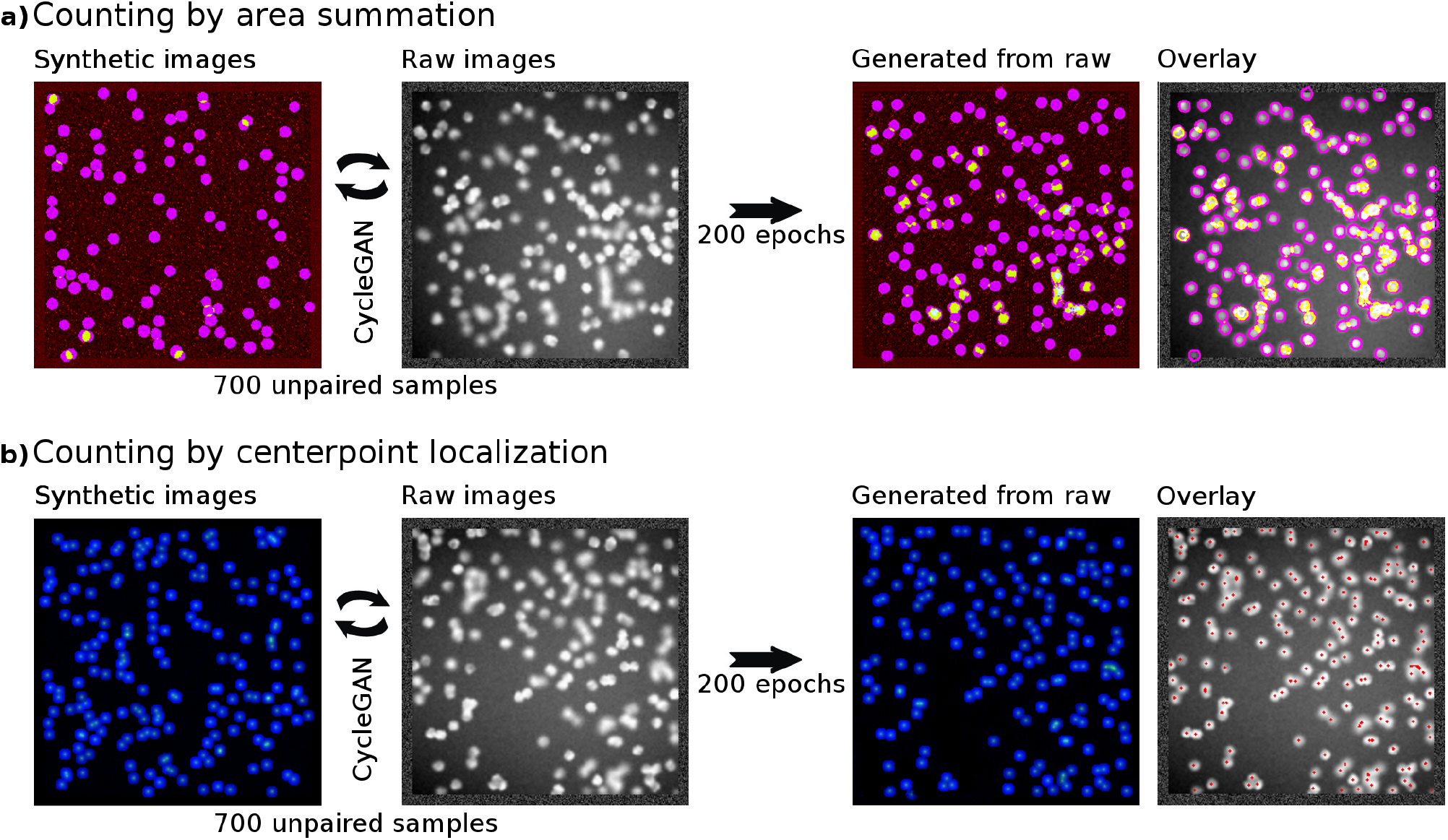
Cell-counting on the VGG dataset: we demonstrate the versatility of UDCT by counting the cells with two distinct datasets. a) A large number of cells, randomly chosen uniformly between 70 and 300, with a radius of 5.5 pixels are placed in a 256×256 image. The background contains gaussian noise in the red channel. The cells are given a single color, and the overlaps are colored distinctly. After convergence, the raw images are transformed and overlaps are weighted in function of how many cells are intersecting in a given area. The background is set to a value of 0. Finally, the image is summed and divided by the cell area to provide a count. b) Here the background has weak noise, the cells are of varying uniform intensity in the blue channel. The green and red channels contain gaussian maps at the cell center, with a standard deviation of 3 in the green and 1 in the red channel. The blue channel is simplifing the task of creating the mapping between the synthetic and the original cells, and the gaussian maps serve the purpose of locating the cell center. These locations are found in the transformed images by maximum likelihood estimation. Finally, in addition to counting the cells by collecting all the center points, we can also retrieve their spatial location.

**Table 1:** Summary on the VGG dataset: UDCT performs slightly worse than Count-ception[19] regardless of which synthetic dataset we use (See Fig.2 a) or b)). However our method is unsupervised and does not directly minimize the mean absolute error during training. Furthermore, we can extract additional information depending on the structure of our synthetic dataset, such as the localization of the cells, which we can pinpoint to within 2 pixels from their ground-truth position (see SI).

|  | Count-ception | UDCT - area summation | UDCT - localization |
| --- | --- | --- | --- |
| Average absolute error | $2.3 \pm 0.4$ | 6.0 | 7.9 |
| Average relative error [%] | - | 3.2% | 4.6 % |

Our approach yields a count accuracy of 96.7% when the synthetic data is setup to perform the counting by area summation (Fig.2a)) and 95.3% when performing the same task by locating the cell centers (Fig.2b)). In the latter case we locate the cells to within 2 pixels of their ground-truth position. This is slightly worse than the state-of-the-art supervised approach of Count-ception. This is to be expected. First, we operate in an unsupervised fashion and do not train our network on ground-truth labels. Secondly, the supervised approach minimizes an objective function that is precisely the counting accuracy (mean absolute error). Our cycleGAN minimizes in qualitative terms a style-transfer accuracy, and in this case indirectly the counting accuracy by inserting information into the transformed data that enables us to count in a post-processing step. Lastly, the VGG Cells dataset contains ground-truth labels from which cells are computer-generated. These labels can in some cases be within a few pixels of one another and generate cells that still look like a single one to the human eye, but correspond to two or more counts based on the ground-truth (see Fig.S2). These are typically cells that are miscounted by our localization approach. However, the likelihood of such cell clusters given a certain number of points randomly distributed in 256×256 space is well-defined and can easily be learned by the supervised network which can correct for them by introducing a systematic bias on its count prediction to minimize the counting error (see Fig.S2). If we correct our localization result by this factor, we bring down the mean absolute error from 7.9 to 4.4. Details of the post-processing steps are given in the Supplementary Information.

With these results we show that the approach is competitive in terms of accuracy, but also much more adaptable and more powerful in terms of the information it can yield. To further demonstrate the power and versatility of the approach, in the next section we localize and count alive and dead cortical primary rat neurons in brightfield microscopy images with high cell density variability and inconsistent image focus.

### Cultures of primary cortical rat neurons

In order to test our network on data that capture the complexity of typical cell images in biomedical research, we created our own dataset of primary cortical rat neurons forming a network on a glass substrate with typical surface-functionalization and cell density. The cells were seeded with a density of 15k/cm^2^ and 50k/cm^2^ onto a poly-(D)-lysine coated glass-slide. The brightfield images have been taken at DIV3 with a 20X objective (Olympus UPLFLN20XPH) that had a numerical aperature of 0.5 (Olympus FluoView 3000 CLSM). Eleven images with a seeding density of 15k/cm^2^ and 16 images with a seeding density of 50k/cm^2^ have been used, each spanning a field-of-view of 1mm^2^. From each image, gradient shifts in the background intensity have been fitted by a third-order 2D polynomial and subtracted. Each 1024 × 1024 pixel image was divided into sixteen 256 × 256 pixel images. All four rotations of each image have been added to the genuine raw dataset for a total of 1728 images. Adding rotated versions of images to the dataset increases the overall size of the dataset, since the network has difficulties recognizing the rotation and subsequently interprets the image as a previously unknown instance. Usually these grayscale (brightfield) images consist of flat-looking, large live neurons and small, round dead neurons. Additionally axons interweave the live neurons creating sharp meandering structures, as can be seen in the top-most row of Fig.3a). A routine characterization of these cultures is performed by counting the amount of live and dead cells in the culture. This is done with fluorescent labels of different colors that selectively target alive and dead cells. Cell labeling can be undesirable as the long-term effects of such stainings on the culture integrity are poorly understood and a successful penetration of such dyes into a cell is not guaranteed. As a consequence, these unpenetrated cells are not labelled and can therefore not be detected using fluorescence.

**Figure 3:**
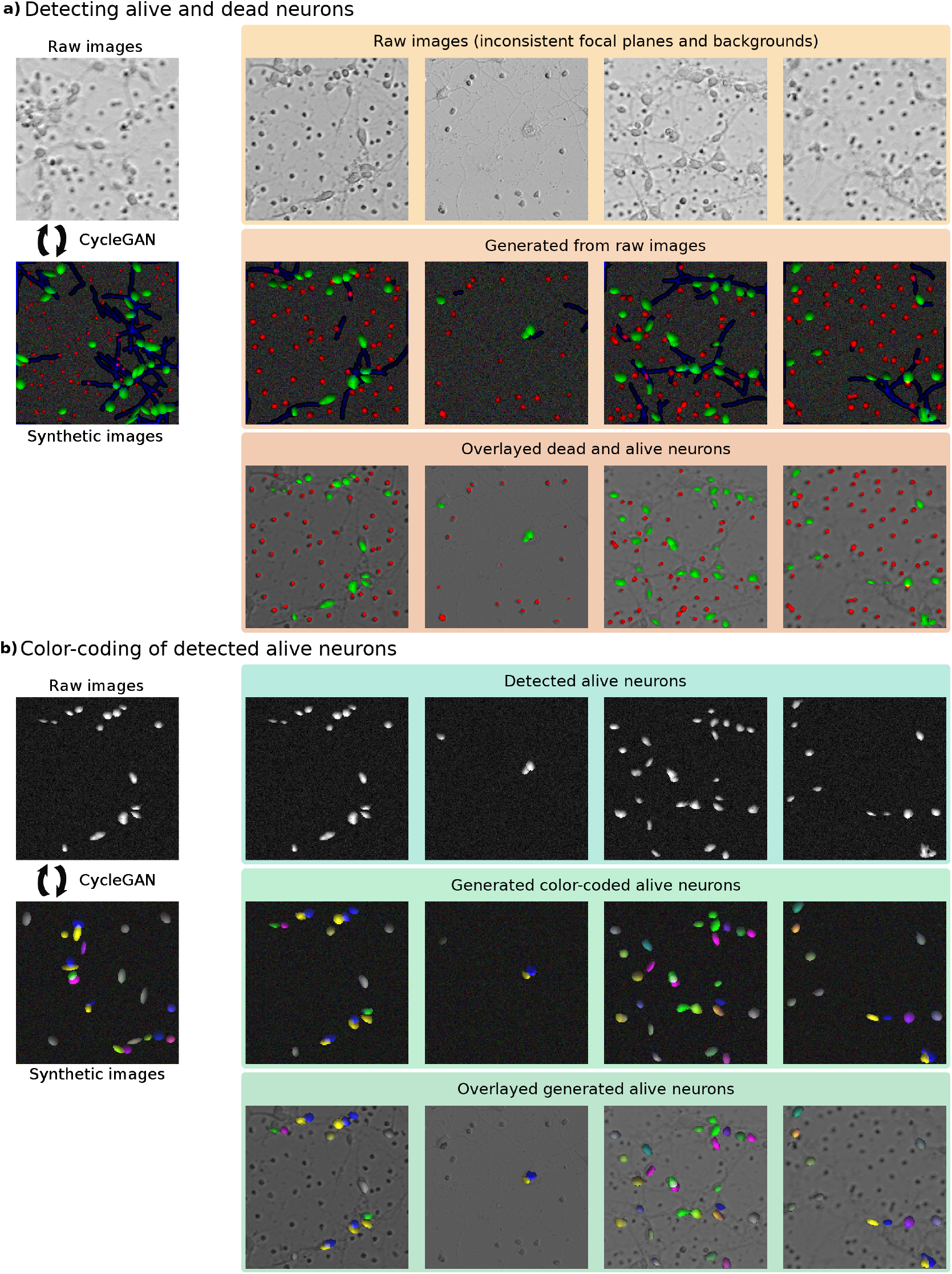
Counting primary rat cortical neurons: We demonstrate the discriminatory power of the cycleGAN by counting the number of alive and dead neurons in a set of images with a wide range of focal blur and backgrounds (top-most row in subfigure a). The dataset is segmented with two cycleGANs. a) The first cycleGAN is trained on synthetic images that consist of small red ellipses and green ellipses. Furthermore, some blue lines have been added to mimic neuritic growth. In the second row, the generated synthetic image out of the raw images shown in the first row are plotted after training for 200 epochs. The third row is an overlap of the predicted dead and alive neurons and the raw images. b) While the number of dead (red) neurons can be easily counted with standard clustering approaches, the alive neurons can overlap, which makes counting difficult. Therefore, a second cycleGAN has been trained on the predictions of the alive cells of the first cycleGAN (top row). Here, the synthetic dataset consists of ellipses with different colors. The colors depend on the location of each ellipse with respect to close-by ellipses. A detailed explanation of the color-coding is given in the supplementary information. The generated color-coded versions of the alive images can be seen in the middle row. The overlayed color-coded predictions on top of the raw images is plotted in the bottom row.

We perform this measurement without the need for fluorescent markers by creating a synthetic dataset containing large green ellipses and small red circles, some blue lines and background noise. To create this syntetic dataset, only a few lines of codes were needed. The details can be found under Dataset construction. In a first step, the cycleGAN maps the raw gray-scale images onto a color-segmented version, whereby the true live and dead cells have been labeled according to the color scheme of the synthetic data. As can be observed in the last row of Fig.3a, counting the dead cells is simple as they do not overlap in most cases. In contrast, live neurons tend to pull together and therefore produce green clusters of pixels larger than a typical cell. Because the focus is on counting, we overcame this issue by creating another synthetic dataset where objects were color-labeled according to their spatial proximity (see Methods, color-coding). The generated images depicted in Fig.3a were thresholded so as to extract only the green (live) pixels and a cycleGAN was performed with proximity-color-coded synthetic data shown in Fig.3b. This allowed to separate clusters of overlapping cells into individual entities (last two rows of Figure 3b). Finally, a simple clustering scheme (see Methods) yielded the live cell count.

We benchmarked our unsupervised counting procedure with the supervised, state-of-the-art Count-ception network by Cohen et. al [19]. To that end, three neural culture experts labeled live and dead cells on 114 images by hand. The counts from different experts is quite variable. The average count per image for each expert is given for both dead and alive neurons in the supplementary Table S1. On average, Expert 3 counts a little over two dead neurons more in each image than Expert 1. The per image count is given in Table 2. The counts for each expert disagrees on average by about 2.3 dead and 1.9 alive neurons. The corresponding mean relative error, which describes by how much the ground truth and the prediction differ, is 5.9 % for the dead neuron count and 10.6 % for the alive neurons. By segmenting the raw images with our cygleGAN networks and applying a simple clustering algorithm (see SI), we can get similar results to the state-of-the-art, even though we achieve this in an unsupervised manner. When counting the alive neurons, we slightly outperform Count-ception.

**Table 2:** Prediction error for neurons: We compare the mean relative error of both the count-ception network as proposed by Cohen et al. [19] and our cycleGAN. The first two rows show the average count and the corresponding standard deviation of both dead and live neurons for each of the three experts. Below, the expert’s ground-truths are compared to each other. The bottom part shows three different prediction methods and their relative error: the error when predicting the average count, our approach, and using the count-ception network proposed by Cohen et al. [19]. When using the average as a network prediction, we get the biggest relative error. This value describes how well any network can perform when basing a prediction solely on a-priori knowledge. For the Count-ception prediction, the networks were trained 5 times. Only the median value is shown, since the network did not always converge (no convergence for at most 1 repetition per point).

| Dead count |  |  |  |
| --- | --- | --- | --- |
|  | Expert 1 | Expert 2 | Expert 3 |
| Average count | 33.4 | 34.4 | 35.9 |
| Std in count | 19.5 | 19.9 | 21.2 |
| Expert 1 | - | $5.4 \pm 5.2 \%$ | $6.6 \pm 5.7 \%$ |
| Expert 2 | $5.4 \pm 5.2 \%$ | - | $5.8 \pm 5.0 \%$ |
| Expert 3 | $6.6 \pm 5.7 \%$ | $5.8 \pm 5.0 \%$ | - |
| Avg. difference to mean | $44.0 \pm 21.8 \%$ | $44.6 \pm 20.8 \%$ | $46.6 \pm 19.2 \%$ |
| Our approach | **$14.1 \pm 13.5 \%$** | **$14.1 \pm 13.1 \%$** | $15.8 \pm 13.2 \%$ |
| Count-ception | $15.7 \pm 12.5 \%$ | $16.4 \pm 13.2 \%$ | **$14.0 \pm 12.4 \%$** |

Table 2: **Prediction error for neurons:** We compare the mean relative error of both the count-ception network as proposed by Cohen et al.
| Alive count |  |  |  |
| --- | --- | --- | --- |
|  | Expert 1 | Expert 2 | Expert 3 |
| Average count | 11.7 | 11.3 | 11.8 |
| Std in count | 8.4 | 8.1 | 8.3 |
| Expert 1 | - | $9.9 \pm 12.9 \%$ | $10.4 \pm 14.7 \%$ |
| Expert 2 | $9.9 \pm 12.9 \%$ | - | $11.5 \pm 16.2 \%$ |
| Expert 3 | $10.4 \pm 14.7 \%$ | $11.5 \pm 16.2 \%$ | - |
| Avg. difference to mean | $48.1 \pm 26.57 \%$ | $46.04 \pm 27.92 \%$ | $47.12 \pm 27.92 \%$ |
| Our approach | **$24.5 \pm 24.4 \%$** | **$25.0 \pm 24.7 \%$** | **$22.2 \pm 23.0 \%$** |
| Count-ception | $39.5 \pm 23.4 \%$ | $36.1 \pm 21.3 \%$ | $49.6 \pm 22.5 \%$ |

### Live-dead assay of C. Elegans

With this dataset we demonstrate the capability of a cycleGAN to distinguish between for- and background in cases where the background has a strong spatial structure. The dataset consists of images of C.elegans exposed to the pathogen *Enterococcus faecalis*. Some of the worms are treated with ampicillin. We investigated, the predictive capabilities of the cycleGAN when it comes to detecting alive (predominately curved) or dead (predominately straight) worms.

The c. elegans infection live/dead image dataset (version 1) was used for the genuine raw images [20]. It was provided by Frederick Ausubel and is available from the Broad Bioimage Benchmark Collection [21]. We discarded the second channel containing the fluorescent images of the GFP labeled dead worms. The dataset consists of 76 grayscale brightfield images with a resolution of 8 bit. These are cropped to 430 × 430 pixels around the field of view. The worms in the images are illuminated with a non-uniform light source, where the point of highest illumination is close to the center region of each image. Besides the worms, the images display small spots of dirt on the surface. For training the network, 750 randomly located 256 × 256 pixel images were cut out of the original images with different rotations (0°, 90^¤^,180^¤^, or 270°) randomly mirrored. Additionally, 750 synthetic images were created for training purposes that also were 256 × 256 pixels in size, and consisted of 3 channels (rgb). The blue channel contains a randomly located 2D gaussian map that mimicks the background. Furthermore, d bright ellipses (major axis: ≈ 6.7, minor axis: ≈ 4.3) have been added to the blue channel, where 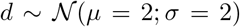, while the numbers of worms added was also guassian distributed *μ* = 7.5 and *σ* = 2 (rounded to the closest integer). The ellipses serve the purpose of mapping dirt contained in the images. Each worm was created with a 50% probability of being alive. A dead worm was created as a predominantly straight line, while the alive one was made curvy. Both are created by a random walk with momentum for the curvature. However, the dead worm is more effected by the momentum causing it to have an overall change in direction that is smaller by a factor of seven in comparison to a live worm. The thickness of the worm was chosen to be 6 pixels with small variations 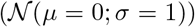 and thinned out ends. The overall length of the worm was set to 125 pixel with a gaussian standard deviation of 15 pixel. These values are based on length measurements done on a small number of worms. The red and green channels encode the local angle *φ* of the worm using [cos(2 · *φ*)/2 + 1, sin(2 · *φ*)/2 + 1]. The results of a modified dataset, where the total number of worms is less (*μ* = 3.5 and *σ* = 0.5) are shown in Fig.S4 for comparison.

We show some segmentation results in Fig.4. The generated synthetic images created by the cycleGAN are used to create an instance segmentation of the individual worms. The segmentation approach is similar to the one described in [22]. In order to predict the viability of the worms, we transformed the colors back to angles and calculated the mean change of the smoothed angle over the length of the whole worm. A two component gaussian mixture model was fitted on the data. All worms that had a higher likelihood of belonging to the gaussian with the higher mean were classified as alive, while the other worms were classified as dead (last row in Fig.4). The only parameter in this classification approach is the standard deviation of the gaussian blur applied on the angle. It was chosen arbitrarily by the authors to be 10. It has not been adapted to minimize the prediction error. A huge effect of the parameter choice on the prediction is not expected. The goodness of prediction was compared to ground truths, for which we marked the center of each curved (assumed alive) and straight (assumed dead) worm. If a worm predicted by the instance classification aligned with such a mark, the worm was assigned this mark as its true label. In total, we were able to find 497 worms that received a dead label and 423 worms with a live label. We compare these labels with the unsupervised approach described above, where a gaussian mixture model was used to predict the type of worm. The confusion matrix for the 920 worms with known label is given in Table (3).

**Figure 4:**
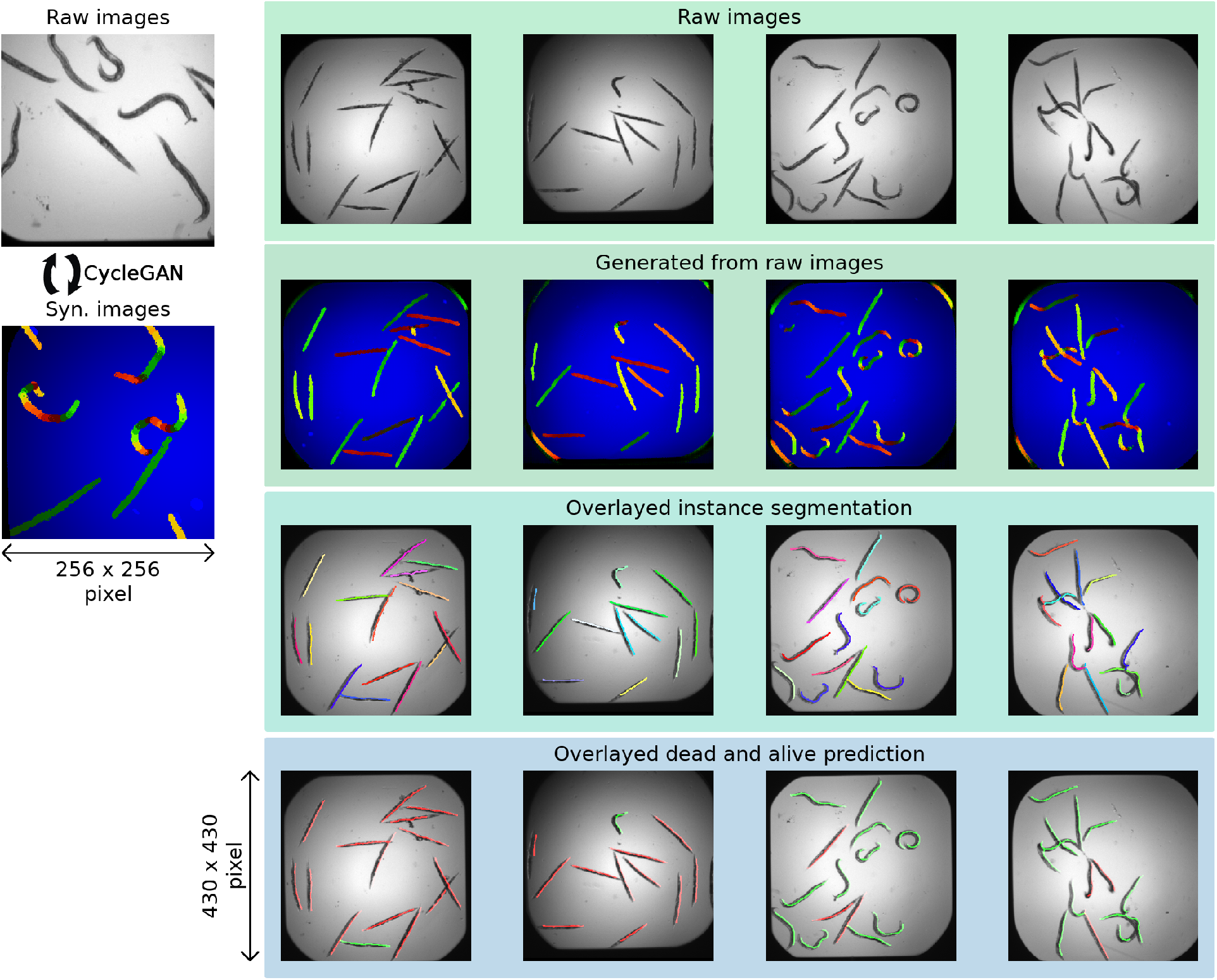
C.elegans viability segmentation: An example cut-out of a raw image used for training the cycleGAN is shown on the left-hand side. On the right-hand side four examples of C.elegans segmentation and classification are given. The first row shows the raw images. In the next row, the corresponding generated segmentation of the cycleGAN is given. Next is the instance segmentation of the worms overlayed over the raw images. In the last row the alive (green) and dead (red) predictions are given.

**Table 3:** Confusion matrix of C. elegans viability: Number of worms for which the viability was correct and incorrect for both alive and dead worms. In brackets, the percentage for predicting a given alive/dead worm (in)correctly is presented.

| 920 worms | Prediction:<br>alive | Prediction:<br>dead |
| --- | --- | --- |
| Ground truth: alive | 399 [94.3 %] | 24 [5.7 %] |
| Ground truth: dead | 68 [13.7 %] | 429 [86.3 %] |

### X-ray computed tomography of silver nanowires

In this section, we show an instance segmentation of fibers (metallic nanowires) in 3D from stacks of 2D images. We acquired high-resolution, X-ray tomography datasets of silver nanowires embedded in polydimeythilsiloxane (PDMS) with the full field Transmission X-ray Microscope of the TOMCAT Beamline at the Swiss Light Source (Paul Scherrer Institute) [23, 24]. This is a common composite material for strain-sensing or stretchable electronics applications. The acquisition volume is 70 micrometers in each direction with a voxel size of 65 nanometers. The nanowire mesh spans the width and height of the volume but is contained in a thickness of 5 microns and is therefore quite flat. Because of this, it is enough to analyze the images along the depth (Fig.5 left). The nanowires are typically 10 micrometers long and vary log-normally in their diameter around an average of 300 nanometers. As seen in Fig.5b, the noise level (produced by the embedding PDMS matrix) is commensurate to the intensity of certain faint wires. The goal is to achieve an instance segmentation of the wire network, meaning that we do not seek to only find where wires are but also to label them individually. To that end, the original dataset, 90 grayscale images of 919 by 832 pixels, was randomly cut into 1000 sub regions 256 by 256 pixels in size. A brief inspection of the cutouts yielded a rough estimate of 10 to 25 wires per image. Figure 5a shows examples of the raw and synthetic images in the dataset used for the cycleGAN. The number of wires is drawn from a uniform distribution between 10 and 25. The position of the wires and their rotation are drawn from a uniform distribution. The wires are between 4 and 10 pixels wide, 30 to 150 pixels long (representing a true width between 150 and 600 nanometers and lengths of 2 to 10 micrometers). They are colored according to the HSV (Hue Saturation Value) color scheme with their orientation coded in the hue (H) and the saturation and value (S,V) set to 1. The background consists of faint gaussian noise. A thousand such pictures were generated. After the cycleGAN converges, the 2D images can be analyzed one by one. The transformed (generated) images are overlaid for qualitative inspection as in Fig.5b. The algorithm works remarkably well and barely misses a wire in the mesh. In order to get a glimpse of the full 3D stack, Fig.5c shows the ray-traced reconstruction of the nanowire mesh in its raw (left) transformed (middle) and instance segmented (right) form. The transformed stack shows that the color coding is consistent from image to image. Because the wires are color-coded with respect to their angle, the pixels can be clustered together into individual wires depending on their hue value, which yields the instance segmentation on the right panel of Fig.5c. This unsupervised ‘one-shot’ clustering of fibers is unique to our knowledge. A previous attempt is described in [7] but it is a supervised training with a heavy post-processing step to disentangle the wire clusters. Because there isn’t any publicly available hand-annotated dataset, we chose not to benchmark our approach. The qualitative results displayed in Fig.5 are convincing in that most of the wires are detected. Furthermore, those samples were produced with 3mg of silver nanowires on a 2-inch circular silicon wafer. This means that we expect about 1080 wires to be located in the field-of-view. The instance segmentation visible on Fig.5c yields 1210 wires.

**Figure 5:**
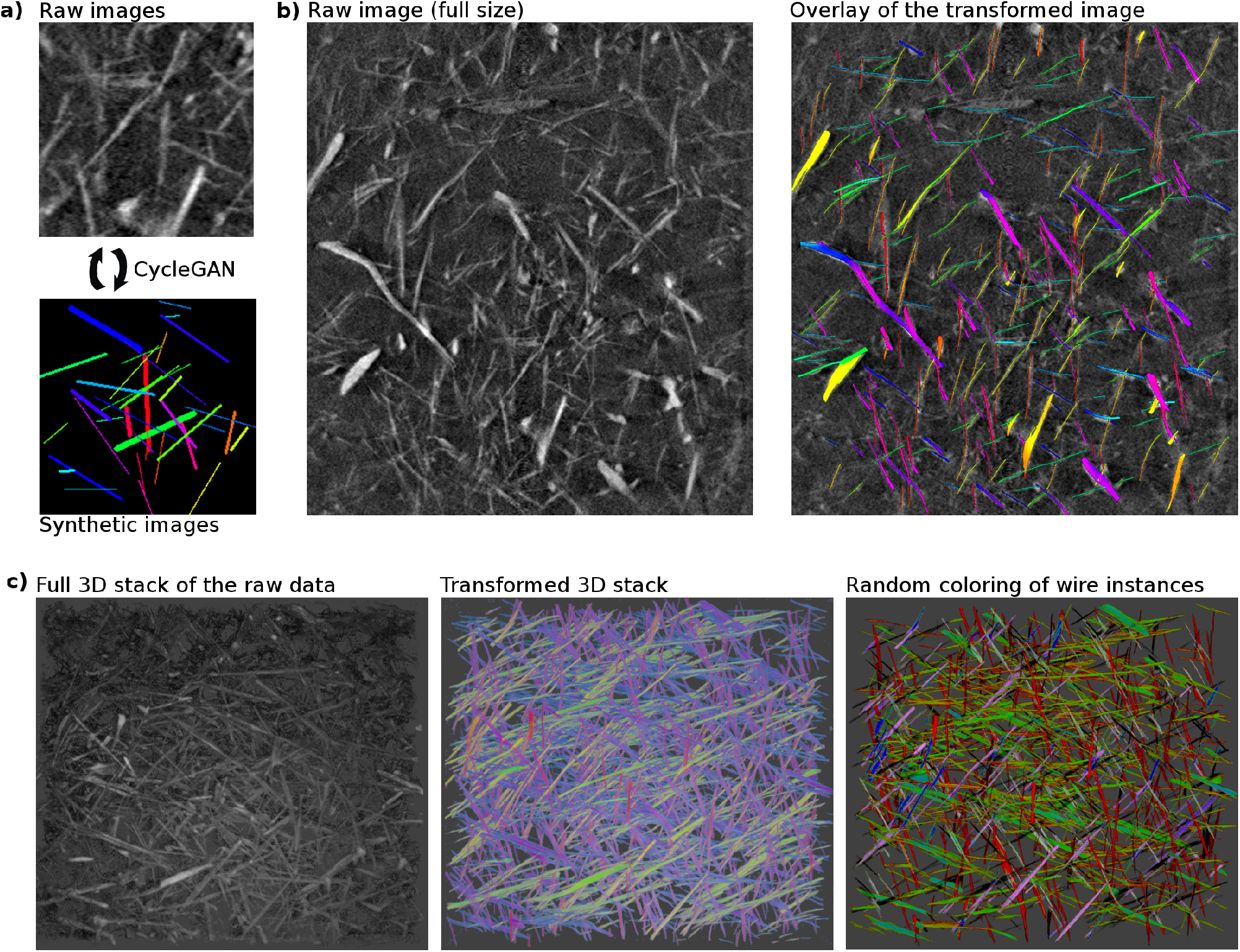
a) Raw and synthetic images used for the cycleGAN to learn the style transfer. Here, the purpose is to label wires in the raw image by color-coding them depending on their orientation. Because the mesh is relatively thin and wires lie flat inside it, one angle is enough to describe them. b) The full images sampled along the stack depth are transformed by the raw-to-synthetic generator. The result is overlaid on a sample image. Most of the wires are detected and labeled correctly. The variation in color of some wires comes from the fact that while the hue is constant, the saturation and brightness are slightly modifed by the cycleGAN. c) 3D-rendering via ray-tracing (Blender) *Left:* Top-view of the raw data displaying a thin mesh of nanowires, reconstructed via ray-tracing where any pixel value lower than a threshold (the noise level) was set to transparent to provide vision into the mesh. *Middle:* same procedure with the transformed images generated by the cycleGAN. The colors are consistent from image to image (into the depth) showing that full wires can be reconstructed by looking at pixel values having the same hue. *Right:* Instance segmentation where adjacent pixel clusters of identical hue were marked with a random color. Small wires are not displayed for clarity. It is clear that these represent the wires and therefore this approach yields a ‘one-shot’ instance segmentation.

The key message of this result is that regular (here, straight wires) data can be instance segmented by creating a very simple synthetic dataset, which in our case took a human less than one hour to implement, instead of hundreds of hours to create training labels for a supervised approach.

## Discussion

We presented an unsupervised approach to segmenting scientific images by creating simple synthetic datasets which encode the information one seeks to extract and letting a cyclic generative adversarial network perform the style transfer between the synthetic and raw data - we call this unsupervised data to content transformation (UDCT). We first showed the power of the approach by performing similarly to the supervised state-of-the-art Count-ception [19] network on a cell-counting task in the VGG Cells dataset. Because we minimize a style-transfer between the raw and synthetic images instead of just counting accuracy, we have the flexibility to encode more information into our synthetic data which in this case yields additionally the location of each cell, a task Count-ception cannot do. This flexibility allowed us to use the same network but with different synthetic data to perform live-dead cell counting in highly variable, inconsistent focus brightfield images of primary rat cortical neurons. We achieved a similar task on an assay of C.elegans with non-trivial background lighting. Finally, we could create a direct instance segmentation of a mesh of nanowires with low signal-to-noise ratio. The latter would be prohibitively time consuming to tackle with standard supervised techniques because of the need for labels during training. Instead, we spent one hour to code the synthetic dataset.

The power of the technique resides in the fact that the researcher can use his or her full creativity to encode the wanted features in a rather simple synthetic data and use the same network architecture to perform the style transfer. This is a direct consequence of the creative power of generative adversarial networks.

The limits of the technique are unclear yet. We propose heuristic observations, such as including a noise and variation component in the objects one wishes to use for segmentation. Concretely, when searching for cells in an image, the synthetic data should have ellipses with varying axes lengths, pixel intensity with some noise, so that the network has a large enough parameter space to perform the mapping between synthetic and raw data. However, there may be cases in which the data is so complex that many equivalent synthetic-to-raw mappings may exist, and finding the right one may be problematic.

In conclusion, we give the scientific community access to an easy-to-use, yet powerful technique to segment complex data. Indeed, one only needs to install the required public software, download our repository and feed into it the raw and synthetic datasets one wishes to analyze. Hopefully this will allow the scientific community to broadly try out this approach and better explore its limits and thereby enhance it. It is our opinion that we need more publicly available challenging biomedical datasets to explore the technique. As much as there is a scientific value in the commonly observed effort to outperform benchmarks by as much as a few percents on public synthetic datasets, the same effort could be spent where there is a wealth of highly complex, unsegmented and biomedically relevant data, where a pragmatic, qualitatively good segmentation would already be satisfactory.

## Methods

### Network architecture

The network architecture is similar to the cycleGAN introduced by Zhu et al. [8] except for three main differences: First, the identity loss has been removed, second, the number of channels has been adapted to the number of channels of the dataset, and third, a novel type of discriminator has been introduced that is discriminating between generated and genuine images using only information that is in the histogram of the image. We call this discriminator the histogram discriminator. The generators are deep convolutional neural networks (CNN) with a similar shape as the SegNet architecture [2]. The deepest encoding level consists of residual layers [25]. To speed up the training of the generator, layer-wise instance normalization [26] was used. The discriminators used a PatchGAN architecture as introduced by Isola et al. [27]. The receptive field of the PatchGAN had a size of 70 × 60. The authors experimented also with 34 × 34 and 142 × 142 PatchGAN as well as a combination of them but could not find qualitative differences.

#### Removing the identity loss

The identity loss according to Zhu et al. preserves the tint between the input and output image of the generators. While this makes sense when transforming Monet’s paintings to photographs, for our tasks it does not. The applications presented in this work rely on encoding features and information into colors which are to be mapped onto various data, be it colored or grayscale. Therefore preserving the tint is not favorable.

#### Adaptive number of image channels

We have noticed that the cycleGAN could often create a perfect cycle reconstruction of a 3-channel image, when it was being mapped on a gray image with 3 channels (all channels have identical values) and back to its original domain. At the same time the GAN responsible for discriminating the synthetic images was causing a mode collapse, a common issue with GANs [28]. We believe that a collapsed image with perfect cycle reconstruction is only possible if the generators ‘‘smuggle” the raw images into the collapsed image (e.g. by putting a compressed version of the image into the collapsed generated image). To circumvent this behavior, we introduced a bottleneck by reducing the number of channels of the gray images to 1, as is often the case for raw data. This measure removed the problem of mode collapse in the GAN responsible for generating raw-looking images. We also believe that this improves the quality of the mappings found by the cycleGAN, as it has much less freedom in the raw-looking images since there’s only 1 channel instead of 3.

#### Introducing a novel histogram discriminator

For the tasks proposed in this work it is critical that the colors created by the generators are identical to their original (synthetic) counterparts in order to simplify segmentation and instance counting. By looking at the histograms of an image, Risser et al. have shown that style transfer becomes more reliable [29]. The histogram loss introduced there could be added as a loss term to the generator. However, one would need to calculate either the loss relative to the whole dataset histograms, which is computationally expensive, or compare it to a subset of the data. The latter proves to be difficult, if there is a high variability between histograms of one image domain (for example a high variability in the number of cells that have to be mapped onto red ellipses). In this work, we propose to add the histogram loss to the discriminator instead of the generator. By doing so, the distribution of the histograms is learned intrinsically for each image domain by the discriminator. It teaches the generator how to change the generated images such that their histograms fit well into the histogram distribution of the genuine images. The histogram discriminator is a multilayer perceptron with two hidden layers. Instead of giving the perceptron a histogram as input, the inverse cumulative density function was given. Both can be transformed into each other. However, the latter can easily be created by sorting an image by its pixel values and is therefore much easier to implement and to backpropagate through.

### Dataset construction

The cycleGAN learns a mapping between two image domains that are independent of each other. Therefore, it is not necessary to find the ground truth for a subset of the images under observation. Instead, the network finds this mapping in an unsupervised manner. The difficulty with such an unsupervised approach is getting the cycleGAN to adopt a meaningful mapping between the two domains. In the following, we introduce a guideline for generating synthetic datasets.

We developed five guiding rules that help achieve meaningful mappings:

1. **Reducing the domain gap** In general, GANs can transform noise to meaningful realistic images. They can also map any image back to noise. An example for such a mapping can be found for our bright-field neuron dataset in Fig.S5. Furthermore, cycleGANs can find a mapping between two image domains even in cases where the domains are conceptually distant. The network does so by replacing patterns found in one domain with patterns of equal probability in the other domain. We hypothesize that the closer the mapping is to an identity mapping, the easier it is for the network to learn it due to its residual layers. Therefore it is easier for the generators to map rod like shapes rather than ellipses to nanowires while at the same time preserving a one-to-one relationship. The point discussed in this guideline is closely related to the fact that transfer learning becomes less effective the more dissimilar the target domain is from the source domain [30]. Furthermore, information like orientation and relative spatial position will not be as easily accessible. We suggest making simple shapes to represent the objects in the raw images and color-coding properties of interest.
2. **Matching object distribution** The discriminator in a GAN tries to find differences between the raw and synthetic images, including the distribution of the object count in the image. If the number of objects drastically differs in the raw and synthetic data, the generator will be forced to remove objects from the oversampled domain and ‘hallucinate” objects from noise from the undersampled domain, which creates a mapping between noise and features that compromises the generator’s ability to find the desired mapping. Such a behaviour can be seen in Fig.S4. The field of view of the PatchGAN has an influence on how well the discriminator can infer the object count distribution. The smaller the field of view, the less of a problem it is to match the object count, since the number of objects the discriminator sees at once is more variable. Our examples show that a perfect match of distributions is not necessary.
3. **Encoding information** The synthetic data must encode the features of interest. Otherwise, it is unlikely that the cycleGAN will find them just by chance. The more different the objects are encoded with respect to the background, the easier it is to discriminate between them in post-processing steps. In this work, the red green blue spaces of images have been used to code information. However, any number of channels can be used. Different properties can be encoded in different channels. In this work, we encoded orientation, cell viability, curvature, relative spatial location and fore-vs-background.
4. **Non-uniform background** In many segmentation tasks one wants to find objects relative to a background. This can prove difficult in case of our network due to the way a CNN works. Since a CNN is deterministic, it will create the same output every time it sees the same input. Therefore, if the synthetic dataset is mostly made out of a monotone-colored background the generator will create monotone-colored images, as well. As a consequence, the discriminator will not look at fine details since it can easily infer from the background alone, whether the image is a genuine or generated image, rendering in turn the use of a GAN futile. We have circumvented this problem in two different ways. Either, noise has been added to the background or a non-uniform background has been inserted into a different channel in the synthetic image. For the synthetic background, guideline 1 and 2 also hold.
5. **Variety of objects** Similarly, the CNN also produces identical shapes in the generated images, if the objects in the synthetic images are identical (e.g. circles of constant color brightness radius). One can see such an example in Fig.S6. This prevents the cycleGAN from finding meaningful mappings, as it cannot generate randomness on its own. To circumvent this, the synthetic data needs some variability for each feature. The generator can use this variability to map the variability of the objects under observation in the raw data. To that extent, changes in color, intensity, shape, and added noise can be used. We hypothesize that more variability in general is better. Such variability should however always be set up in a way, that features of interest can easily be distinguished during post-processing.

## Code availability

Our tensorflow implementation of the cycleGAN with our novel histogram discriminator, all the synthetic data generating scripts along with the analysis and post-processing scripts are available at https://github.com/UDCTGAN/UDCT.

## Data availability

The brightfield dataset of primary cortical neurons with the ground-truth labels,the annotated C.elegans dataset,the X-ray-computed tomography of silver nanowires is available and all synthetic datasets and trained networks are available at https://downloads.lbb.ethz.ch

## Acknowledgements

We would like to give special thanks to Prof. Dr. Shiva Tyagarajan, Prof. Dr. Jean-Marc Fritschy, and Giovanna Bosshard from the Institute of Pharmacology and Toxicology - Morphological and Behavioral Neuroscience at the University of Zurich for generously providing us with rat hippocampal tissue. We also thank Stephen Wheeler (ETH) for technical support and the Swiss National Science Foundation, ETH Zurich and the FreeNovation grant for financial support. We acknowledge the Paul Scherrer Institut, Villigen, Switzerland for provision of synchrotron radiation beamtime at beamline TOMCAT of the SLS.

