## Supplementary Information for "UDCT: Unsupervised data to content transformation with histogram-matching cycle-consistent generative adversarial networks"

February 28, 2019

[1]Laboratory of Biosensors and Bioelectronics, ETH Zurich, Gloriastrasse 35, 8092 Zurich, Switzerland
[2]Paul Scherrer Inst, Swiss Light Source, CH-5232 Villigen, Switzerland
[3]Institute for Biomedical Engineering, University and ETH Zuurich, Zurich, Switzerland

### Supplementary Information

#### Network formulation

Let $A \in [0,1]^{n\times m\times a}$ be the set of real images with dimensions $n$ by $m$ and $a$ channels. Furthermore, $B \in [0,1]^{n\times m\times b}$ be the set of synthetic images of same size and $b$ channels. The generators $\mathcal{G}_B : A \to B$ and $\mathcal{G}_A : B \to A$ transform a real image into a synthetic images and a synthetic image to a real one, respectively. The discriminator $\mathcal{D}_\mathcal{X} : \mathcal{X} \to [0,1]$ where $\mathcal{X} \in \{A, B\}$ predicts, whether a data sample is either a genuine image (real or synthetic) or a generated image. A genuine image was encoded with a 0, while a generated image is encoded with a 1. The histogram discriminator $\mathcal{H}_\mathcal{X} : \mathbb{R}^h \to [0,1]$ functions as $\mathcal{D}_\mathcal{X}$ with the exception that it will get a binned inverse cumulative distribution function ($\text{iCDF}_h(\cdot)$) of the input image with $h$ bins.

In the following, the network architecture is described. Let `cUsV-W` be a `U` × `U` convolution layer with stride `V` and `W` filters. Futhermore, `IN` denotes an InstanceNorm layer, `ReLU` a rectified linear unit activation, `leaky_ReLU` a leaky rectified linear unit activation with (alpha = 0.2), and `tanh` a hyperbolic tangent activation function. The shorthand for a fractional-strided `U` × `U` convolution layer with stride `1/V` and `W` filters is `ctUsV-W`. A residual addition is signified with `r<`. The layer to which the network is added is marked with the most recent `>` before the residual layer. A fully connected layer with `W` nodes and a dropout probability of 50% is defined as `dW`. Finally, `pU` describes a 2 dimensional reflective padding step.

**Generator** The generators are built in a similar fashion as the generator with 6 residual layers proposed by Zhu et al. [1]. Below, $\mathcal{W}$ describes the number of channels in the output domain ($a$ or $b$).

p3 c7s1-64, IN, ReLU
p1, c3s2-128, IN, ReLU
p1, c3s2-256, IN, ReLU
>, p1, c3s1-256, IN, ReLU, p1, c3s1-256, IN, r< [repeat 6 times]
ct3s2-128, IN, ReLU
ct3s2-64, IN, ReLU
c7s3-$\mathcal{W}$, IN, tanh/2 + 0.5

**Discriminator** Both discriminators are identical to the discriminator introduced in Zhu et al. [1]. The discriminator has a receptive field of size 70 × 70.

p1, c4s2-64, leaky_ReLU
p1, c4s2-128, IN, leaky_ReLU
p1, c4s2-256, IN, leaky_ReLU

`p1`, `c4s1-512`, `IN`, `leaky_ReLU`
`p1`, `c4s1-1`

**Histogram discriminator** The histogram discriminator gets an inverse cumulative distribution function (iCDF) as input. The iCDF can be transformed into a histogram. However, it is easier to create, since it is equivalent to the intensity-sorted pixel values of an image. To reduce the complexity of the network, the iCDF is binned into $h$ by taking the mean of all the elements belonging to the bin. Since $h$ was set to $n$, the number of averaged values for each bin was $m$. If an image had multiple channels, each channel was binned separately from each other. Afterwards, the channel histograms were concatenated and fed into the network.

The histogram discriminator is a multi-layer perceptron with two hidden layers.
`d64`, `tanh`, `d64`, `tanh`, `d1 + 0.5`

### Losses

The total loss of the network can be described by:

$$\mathcal{L}_{\text{total}} = \mathcal{L}_{\text{dis\_A}} + \mathcal{L}_{\text{dis\_B}} + \lambda_c \cdot \mathcal{L}_{\text{cyc}}, \tag{S1}$$

where $\lambda_c$ is an arbitrary value weighting the cycle-consistent loss against the adversarial losses. The terms on the right side are defined below. The optimizer is trying to solve:

$$\mathcal{G}_A^*, \mathcal{G}_B^*, \mathcal{D}_A^*, \mathcal{D}_B^*, \mathcal{H}_A^*, \mathcal{H}_B^* = \underset{\mathcal{G}_A^*, \mathcal{G}_B^*}{\text{argmin}}\, \underset{\mathcal{D}_A^*, \mathcal{D}_B^*}{\text{argmax}}\, \underset{\mathcal{H}_A^*, \mathcal{H}_B^*}{\text{argmax}}\, \mathcal{L}_{\text{total}} \tag{S2}$$

The network is implemented with two optimizers. The first one trains the four discriminators by maximizing equation (S2), while the second one trains the two generators by minimizing equation (S2).

**Adversarial loss** The adversarial loss used in this work is the same as the usual least-square loss for GANs with the addition of the loss function given by the histogram discriminator. The optimizer for the discriminators tries to maximize the adversarial losses. In the equation below, $\mathcal{X}$ is either $A$ or $B$.

$$\begin{aligned} \mathcal{L}_{\text{dis\_}\mathcal{X}} = {} & \mathbb{E}_{x\sim p_{\text{data}}}\left[(1 - \mathcal{D}_\mathcal{X}(x))^2\right] + \mathbb{E}_{y\sim p_{\text{data}}}\left[(\mathcal{D}_\mathcal{X}(\mathcal{G}_\mathcal{X}(y)))^2\right] \\ & + \lambda_h \cdot \left(\mathbb{E}_{x\sim p_{\text{data}}}\left[(1 - \mathcal{H}_\mathcal{X}(x))^2\right] + \mathbb{E}_{y\sim p_{\text{data}}}\left[(\mathcal{H}_\mathcal{X}(\mathcal{G}_\mathcal{X}(y)))^2\right]\right) \end{aligned} \tag{S3}$$

**Cycle-consistent loss** The cycle-consistent loss encourages the generators to approximate a bijective mapping between the relevant subsets of $A$ and $B$. The cycle-consistent loss consists of a loss term for the cycle $A \to B \to A$ and a term for the cycle $B \to A \to B$.

$$\mathcal{L}_{\text{cyc}} = \mathbb{E}_{x\sim p_{\text{data}}}\left[(x - \mathcal{G}_A(\mathcal{G}_B(x)))^2\right] + \mathbb{E}_{y\sim p_{\text{data}}}\left[(y - \mathcal{G}_B(\mathcal{G}_A(y)))^2\right] \tag{S4}$$

**Generator loss** The two generator losses are used to train the generators. The generator optimizer is trying to minimize these losses. The cycle-consistent loss is divided by a factor of two in order to make the overall loss identical to the one given in equation (S1).

$$\mathcal{L}_{\text{gen\_}\mathcal{X}} = \mathbb{E}_{y\sim p_{\text{data}}}\left[(\mathcal{D}_\mathcal{X}(\mathcal{G}_\mathcal{X}(y)))^2\right] + \lambda_h \cdot \mathbb{E}_{y\sim p_{\text{data}}}\left[(\mathcal{H}_\mathcal{X}(\mathcal{G}_\mathcal{X}(y)))^2\right] + \frac{\lambda_c}{2} \cdot \mathcal{L}_{\text{cyc}} \tag{S5}$$

### Training

Unless otherwise stated, each dataset has been trained with a $\lambda_c$ of 10 and a $\lambda_h$ of 0. An optimizer ($\beta_1 = 0.5$, $\beta_2 = 0.999$, $\epsilon = 10^{-8}$) has been used to MinMax equation (S2). Each network has been trained for 200 epochs. The learning rate was fixed to 0.0002 for the first 100 epochs. Afterwards it linearly decayed to 0 for the next 100 epochs. As in Zhu et al. [1], all convolutional kernels were initialized from a gaussian distribution (mean: 0, std: 0.02). In order to achieve a more consistent cyclic behaviour, noise was added to the input of all discriminators. The noise added was gaussian noise with standard deviation of $0.9^n$ where $n$ is the current epoch. In order to follow the implementation of Zhu et al. as close as possible, a buffer was introduced that stored 50 previously generated images on which the discriminator were trained as described by Shrivastava et al. [2]. The batchsize was fixed to 4. The buffer was filled with all 4 images until it was full. Afterwards, four random elements in the buffer were replaced with new ones. Following the suggestion of Shrivastava et al., the discriminators were trained with 2 images from the buffer and 2 current ones.

### VGG-Cells

With the synthetic dataset displayed on Fig.2b), we localize cells to within 2 pixels on average from their ground truth position. The distribution of distance between a detected cell and its closest ground-truth position is displayed on Fig.S1.

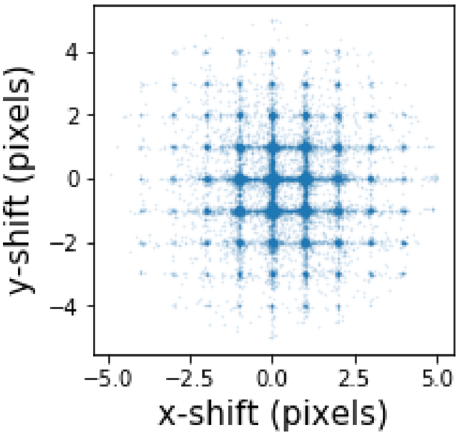


Figure S1: Distance-vectors between detected VGG Cells centers and the ground truth labels in x-y pixel shift units, displaying 34'000 detected cells. Each point displays the x-y shift between a generator cell's position and the nearest ground-truth label. The average error on the location of the cell-center is 1.25 pixels. These numbers are fractional because the maximum likelihood estimation of the centers of the gaussians in the transformed dataset yield real and not integer numbers.

On average we count 95.4% of the cells. Because we operate in an unsupervised fashion, we do not directly minimize the counting error during training. We notice however that some cells would not be counted either by a human, as shown in Fig.S2. Indeed, the ground-truth labels used to create the VGG Cells dataset in some cases can be so close to one another that the generated cell ensemble they represent only looks like a single cell. If a human cannot distinguish these cells, our cycleGAN approach cannot either. However, the number of cell-centers that are within a few pixels of one another shows as expected a pure $N^2$ dependency, where $N$ is the number of cells-centers in an image (see Fig.S2). This means that a supervised network can easily minimize its count error by adding a bias of $aN^2$ to its count $N$, in order to reduce the counting gap. If we take our raw cycleGAN cell count based on the results of Fig.2b, that we define as $c$, for each image and let $c + \alpha c^2$ be a modified count and minimize the error against the true counts, we bring down our mean average error to 4 cells per image.

### Color-coding neuron location

The color of each neuron in the synthetic images of the color-coded neuron dataset only depends on the center location of each neuron. Let $C_p \in \{0,1\}^{n\times m}$ be one, if the pixel describes the center of a neuron in the $p^{\text{th}}$ synthetic image and zero otherwise. Furthermore, let $K \in \mathbb{R}^{101\times101\times3}$ be a kernel used for determining the colors. The neuron in the $p^{\text{th}}$ image with location $(u, v)$ is getting an rgb color vector of

$$\frac{\tanh\left([C_p(\cdot,\cdot) * K(\cdot,\cdot,k)]\,(u,v)\right) + 1}{2}, \tag{S6}$$

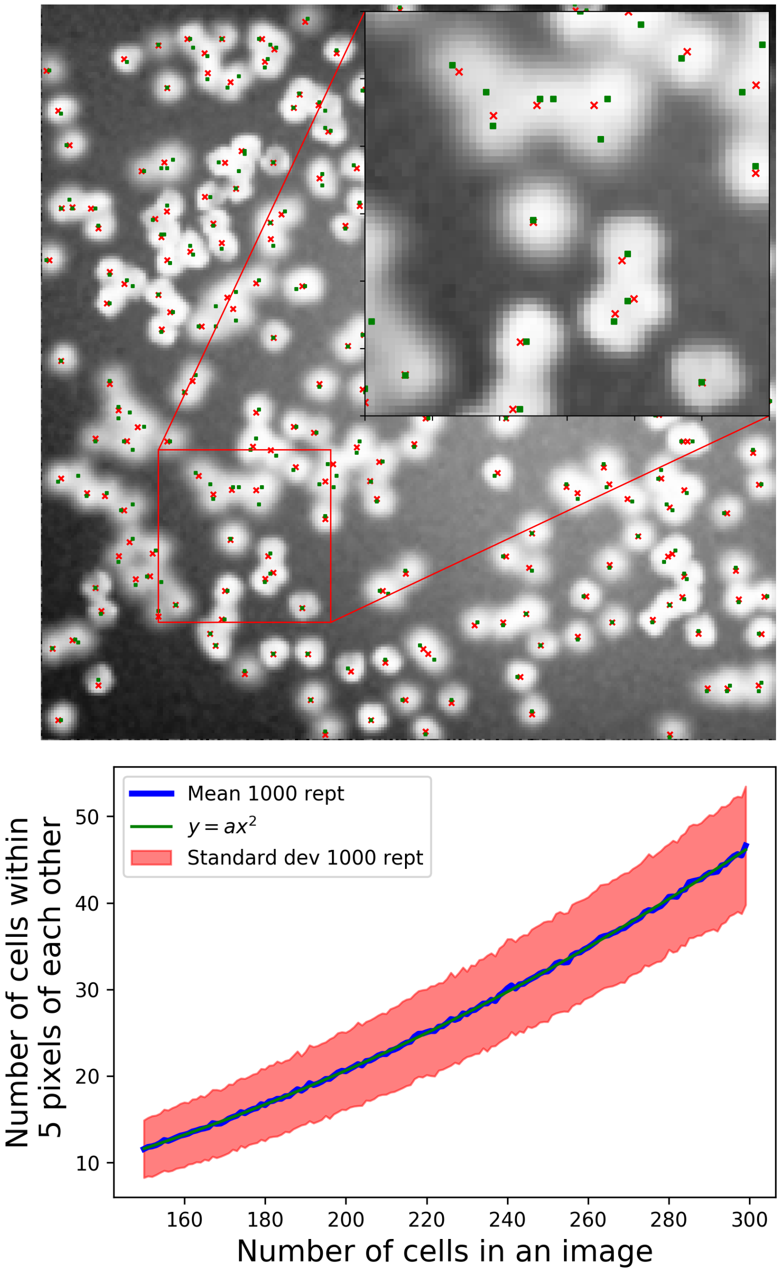


Figure S2: **Issues with VGG cells dataset**: *Top*: the green dots are the ground-truth positions of the cells of the VGG dataset. The red dots are the cell-center locations detected by our cycleGAN method displayed in Fig.2b. The inset shows that in some cases even a human could not count the cells correctly based on the ground-truth positions. This is particularly evident in the top part of the inset. *Bottom*: We distributed $N$ cells-centers randomly in 256×256 pixel space, and counting how many are within $m$ (here 5) pixels of each other and repeated it a 1000 times. The blue curve shows the mean number of such cells, and the red spread is the standard deviation of the 1000 repetitions. The number of such cells follows an $N^2$ relation. This means that when a supervised network minimizes the error count, although it may inherently count the same cells as our cycleGAN, it has the ability to easily correct for the error by adding a bias of $\alpha x^2$, where $x$ is what the network would truly count.

where $k \in \{\text{red}, \text{green}, \text{blue}\}$ describes the channel. The kernel K is shown in Figure (S3) and can be fully described by the matrix $\{\mathbf{K}\}_{i,j}$:

$$K = [\mathbf{K}^{\mathrm{T}}; 1 - \mathbf{K}; \mathbf{K}] \tag{S7}$$

with

$$\mathbf{K}_{i,j} = f(i) \cdot g(j), \qquad i, j \in \{1, 2, \dots, 101\}, \tag{S8}$$

where the functions $f(\cdot)$ and $g(\cdot)$ are defined as

$$f(i) = \begin{cases} \max\left\{ \min\left\{\frac{3.75}{i}, 3\right\}, -3\right\} & \text{if } i \neq 50 \\ 0 & \text{if } i = 50 \end{cases} \tag{S9}$$

and

$$g(j) = \begin{cases} \min\left\{\frac{3.75}{|j|}, 3\right\} & \text{if } j \neq 50 \\ 3 & \text{if } j = 50. \end{cases} \tag{S10}$$

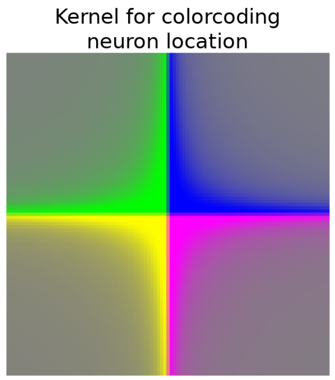


Figure S3: The kernel K used to determine the color of each neuron in the synthetic dataset. The color only depends on the relative location of a neuron with respect to other neurons.

### C.elegans dataset

The number of worms put in the synthetic dataset influences the quality of the cycles. In this case, the synthetic dataset does not have enough worms.

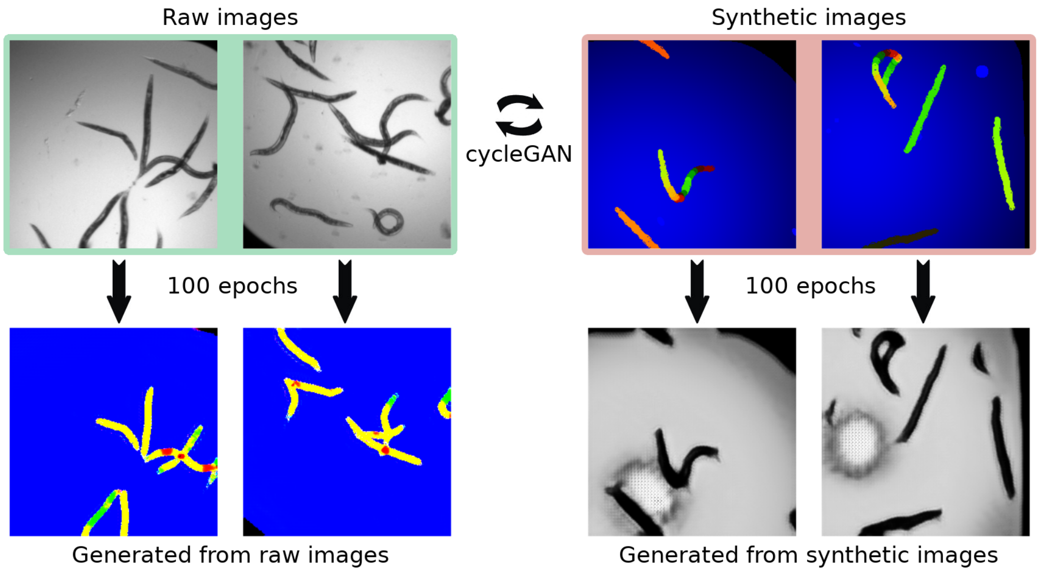


Figure S4: An example synthetic dataset with approximately half the number of worms as in the raw dataset. Especially in raw images with a lot of worms, some worms are removed by the generator (bottom left). The opposite effect occurs in the generator that creates raw images (bottom right), where worms are created out of the background and the image boundary.

### Primary cortical neurons dataset : Example of a backchannel

We illustrate here the issue that can arise without histogram loss in Fig.S5. On the top row, Fig.S5a shows the raw image cycle. Because of the residual network in the generator, part of the image is compressed into the generated image, but not clearly visible at all. However it is perfectly retrieved in the cycle. We call this a backchannel, as the original image is somehow retrieved through the cycle, without having a proper generated image.
This backchannel is much more difficult for the network to achieve in the synthetic cycle shown in Fig.S5b. We hypothesize that this is due to the fact that there is only 1 channel available in the generated (raw domain) image.
The effect of the histogram loss is more thoroughly discussed in Fig.S7.

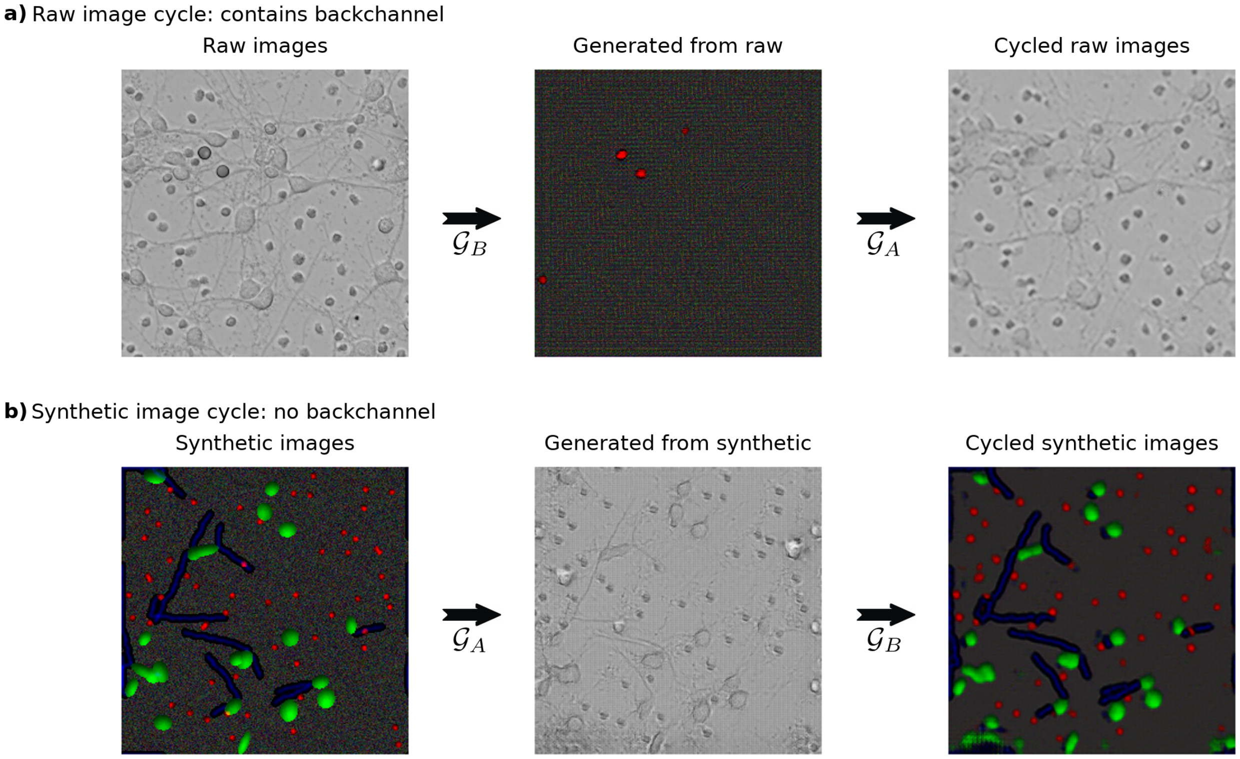


Figure S5: The figure shows backchannel. For the raw image cycle (a), a raw image is mapped to a generated synthetic image without any clear relationship. However, the cycled image is a close match to the original image. At the same time, the synthetic image cycle (b) appears to function as desired.

### Importance of variation in synthetic images

We display the importance of having enough variation in the synthetic dataset to generate realistic raw images in Fig.S6. CNNs and therefore GANs are deterministic functions: for the same input they always generate the same output. Real objects like neurons on a glass slide come in different kinds of shapes, size, and brightness. Therefore, the synthetic ellipses that should represent them in the synthetic data should bear similar amounts of variation. Otherwise, the complexity of real neurons will not be generated as visible on the left side of Fig.S6. On the other hand, by varying slightly the shape, the intensity profile and the background noise, we can create much more realistic pictures. Although we are only interested in the synthetic-looking generated pictures for the analysis, since those are the transformed raw images, it is very important to have a good cycle both in the raw and the synthetic cycle to have a good mapping between synthetic and raw data.

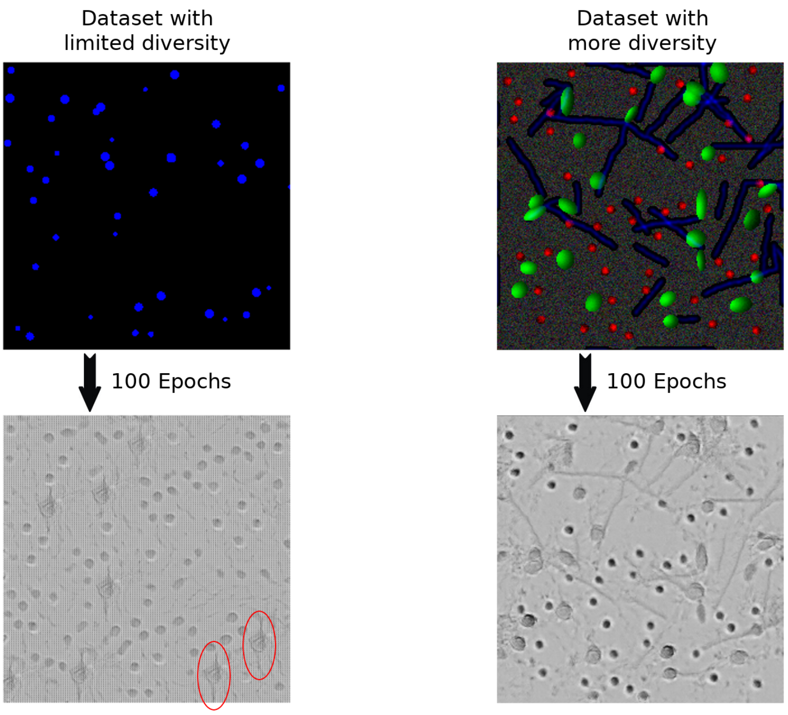


Figure S6: The effect that the diversity of a dataset can have on the generated images. The brightfield neuronal images were trained with a cycleGAN using two different synthetic datasets. When the diversity of the dataset was low (left side), repetitive patterns occur in the generated images (see circled data). When the complexity of the synthetic dataset is higher (right side), the alive neurons can exhibit a wider range of possible shapes. Both networks have been trained for 100 epochs.

### Importance of histogram loss

On Fig.S7 we compare the generated images with and without histogram loss. One can see that quite often, the synthetic-looking generated pictures without histogramm loss (second row of green shaded panel on Fig.S7) are often not good. This goes back to the backchannels discussed in FigS5.

### Mean absolute and relative error

The mean absolute and relative error of the neuronal counts is given in Table (**??**) for the bright-field cortical neurons. The mean absolute error is defined as

$$\text{Error}_{\text{abs}} = \frac{1}{|\Omega|} \sum_{i \in \Omega} |\text{pred}_i - \text{gt}_i|, \tag{S11}$$

where $\Omega$ is the set of all datapoints, $|\Omega|$ the number of elements in $\Omega$, $\text{pred}_i$ the number of predicted neurons for the $i^{\text{th}}$ datapoint, and $\text{gt}_i$ the corresponding groundtruth. In a similar fashion, the mean relative error is defined as

$$\text{Error}_{\text{abs}} = \frac{1}{|\Omega|} \sum_{i \in \Omega} \begin{cases} 1 - \frac{\min\{\text{pred}_i, \text{gt}_i\}}{\max\{\text{pred}_i, \text{gt}_i\}} & \text{if } \max\{\text{pred}_i, \text{gt}_i\} \neq 0 \\ 0 & \text{if } \max\{\text{pred}_i, \text{gt}_i\} = 0. \end{cases} \tag{S12}$$

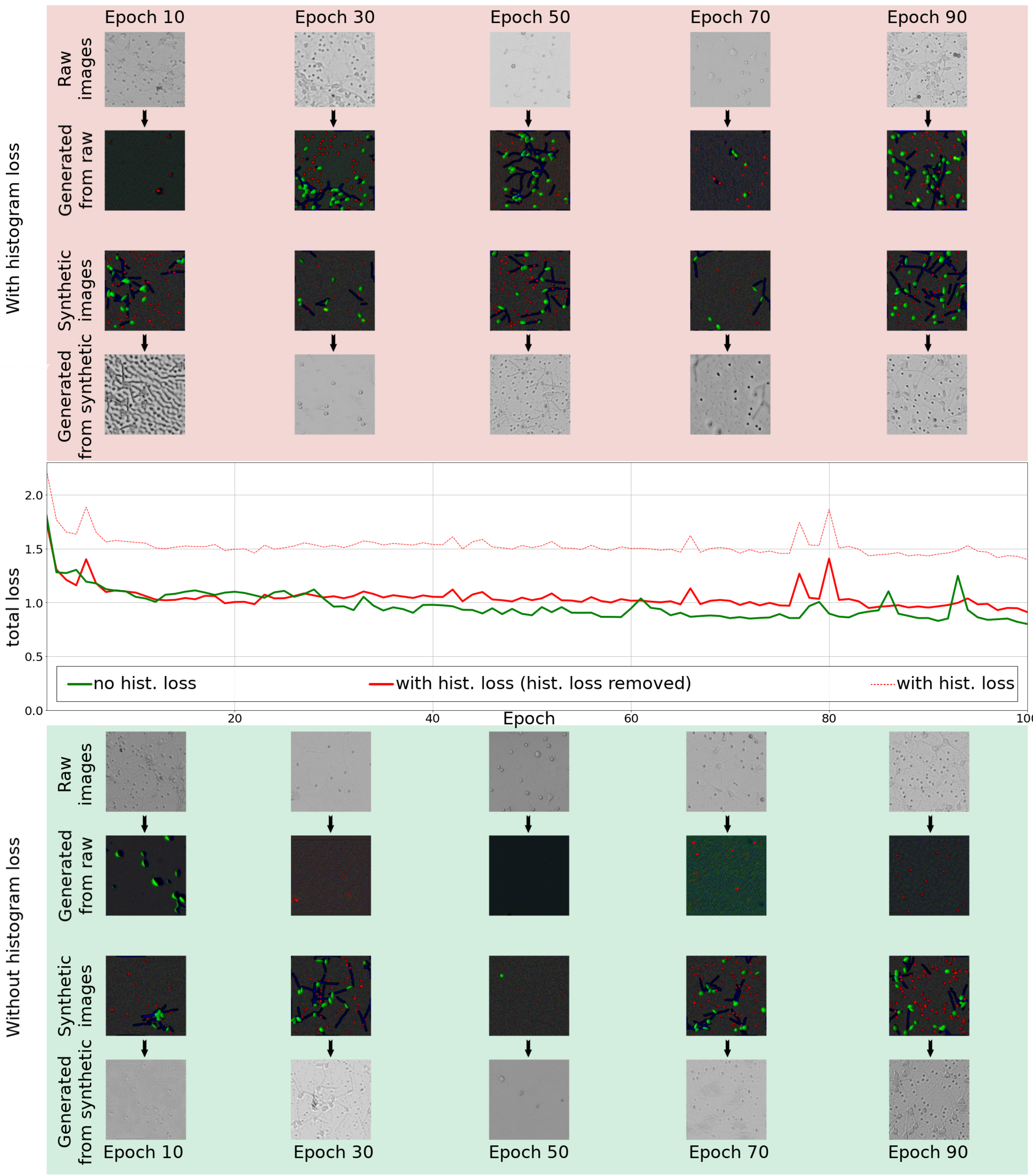


Figure S7: Comparison between having a histogram discriminator versus not having it. The total loss, as defined in Equation S1, is plotted for the first 100 epoch of training in the case where the histogram discriminator is not used (continuous green) and when it is (dashed red line). For better comparison of the two cases, the latter has also been plotted after removing the contributing terms of the histogram discriminator from the total loss (continuous red). Above and below the losses, example generated images are shown for different times during the training when not using a histogram discriminator and when using it, respectively. The authors believe that there are two reasons for the lower total loss when not using a histogram even though the quality of the created images is qualitatively worse. First, the total loss with histogram discriminator is minimized by the network (dashed red) instead of the total loss not considering the histogram losses (continuous red). Second, the lower quality of the generated images helps the discriminators to make better predictions. The corresponding improvement for the loss is bigger than the cost associated with a worse cycle reconstruction.

### Supplementary Tables

| | Expert 1 | Expert 2 | Expert 3 |
|---|---|---|---|
| Average dead count | 33.41 | 34.39 | 35.86 |
| Average alive count | 11.68 | 11.26 | 11.76 |
| Average total count | 45.10 | 45.65 | 47.62 |

Table S1: **Average number of neurons per image**: The average number of dead, alive, and total neurons as counted by three different experts. While the average alive count is quite consistent for each expert, the average number of dead cells counted can be off by as much as 7%.

**Mean absolute error**

| | Dead count | | | Alive count | | |
|---|---|---|---|---|---|---|
| | Expert 1 | Expert 2 | Expert 3 | Expert 1 | Expert 2 | Expert 3 |
| Average count | 33.4 | 34.4 | 35.9 | 11.7 | 11.3 | 11.8 |
| Std in count | 19.5 | 19.9 | 21.2 | 8.4 | 8.1 | 8.3 |
| Expert 1 | - | 1.9 ± 2.1 | 2.7 ± 2.9 | - | 1.4 ± 1.9 | 1.2 ± 1.5 |
| Expert 2 | 1.9 ± 2.1 | - | 2.2 ± 2.4 | 1.4 ± 1.9 | - | 1.1 ± 1.5 |
| Expert 3 | 2.7 ± 2.9 | 2.2 ± 2.4 | - | 1.2 ± 1.5 | 1.1 ± 1.5 | - |
| Predicting avg | 18.1 ± 7.4 | 18.6 ± 7.1 | 20.1 ± 6.8 | 7.2 ± 7.4 | 6.8 ± 7.1 | 7.1 ± 6.8 |
| Our approach | **3.9** ± 3.7 | **4.4** ± 4.4 | 5.4 ± 5.3 | **2.4** ± 2.3 | **2.4** ± 2.6 | **2.3** ± 2.4 |
| Count-ception | 5.1 ± 6.4 | 5.6 ± 5.7 | **4.7** ± 5.3 | 4.4 ± 4.5 | 3.4 ± 3.1 | 5.8 ± 4.7 |

Table S2: **Average number of neurons per image**: The average number of dead, alive, and total neurons as counted by three different experts. While the average alive count is quite consistent for each expert, the average number of dead cells counted can be off by as much as 7%.

### Post-processing

#### VGG Dataset

**Counting by area**: the raw-to-synthetic generated images are the color-labeled versions of the raw images. The images are in a shape of [276,276,3] floats between 0 and 1, where the last 3 channels are the colors. Because the last (blue) channel contains the noise, to convert the colors into numbers, we apply the following transformation to image Im

$$Im[:,:,0] < 0.2 \quad \& \quad Im[:,:,1] > 0.2] \to 1 \tag{S13}$$

$$Im[:,:,0] > 0.2 \quad \& \quad Im[:,:,1] < 0.2] \to 2 \tag{S14}$$

$$Im[:,:,0] > 0.2 \quad \& \quad Im[:,:,1] > 0.2] \to 3 \tag{S15}$$

signifying that we map green regions to 1 (no overlap), red regions to 2 (two cells overlap) and the white regions to 3 (more than 2 overlaps). The other pixels are mapped onto 0. The resulting image, $Im_t$, is now a [276,276] array of integers between 0 (background) and 3. The resulting transformed image is then summed up and divided by the average cell radius $r$ (5.45 pixels) in the synthetic data. Therefore,

$$\text{Count} = \sum_{i,j=1}^{276} \frac{Im_t[i,j]}{\pi r^2} \tag{S16}$$

**Counting by position**

The raw-to-synthetic generated images contain the cell shapes in the red channel, and gaussian maps $G(x, y, x_0, y_0) = e^{-(x-x_0)^2-(y-y_0)^2}$ with a standard deviation of 1 pixel in the blue channel at the center of each cell. We need to find the coordinates $x_0, y_0$ of each gaussian map, without knowing how many there are. To that end, the following algorithm is performed:

1. Threshold the image in the blue channel
2. Find all pixel clusters above the threshold
3. For each cluster, perform a fixed-covariance (variance of 1) gaussian mixture model with a variable number of gaussians. Choose the number that minimizes the distance between the pixel values and the fit
4. Store all the found gaussian centers

Correction factor: for each pixel cluster whose maximal pixel value was above 1, but we could not separate into two distinct gaussians (they are too close), we added a count of 1 to the number of found centers.

#### Primary cortical neurons

**Counting dead neurons:** For predicting the location of dead neurons, the red channel of the cycleGAN generated images were gaussian-smoothed with a mean of half a pixel. A dead neuron was predicted at every peak in the smoothed image that was at least as high as 0.5 (pixels can have values between 0 and 1) and did not have any higher peaks in a distance of 2 pixels.

**Counting live neurons:** The live neurons were predicted by first creating a binary map $B$, which was 1 for each pixel of image $I$, where the sum of the three color channels of the generated images of the second neuron cycleGAN exceeded 0.75. All other pixels were set to 0.

$$B[x, y] = \begin{cases} 1 & \text{if } I_{\text{red}}[x, y] + I_{\text{green}}[x, y] + I_{\text{blue}}[x, y] > 0.75 \\ 0 & \text{otherwise} \end{cases} \tag{S17}$$

From $B$, all connected pixel clusters were extracted. Islands were discarded, if they contained less than 10 pixels (noise). All remaining islands correspond to one or more live neuron.

For each island, we seek to find the number of distinct colors, since this number corresponds to the number of cells for the given island. Below calculations are done for each island separately. To simplify readability, the below equations do not contain an index for each island. To find the number of live neurons, we first normalized the color vector of each pixel belonging to the island under observation to one.

$$\text{Col}_i = \frac{1}{\sqrt{I^2_{\text{red}}[x_i, y_i] + I^2_{\text{green}}[x_i, y_i] + I^2_{\text{blue}}[x_i, y_i]}} \begin{bmatrix} I_{\text{red}}[x_i, y_i] \\ I_{\text{green}}[x_i, y_i] \\ I_{\text{blue}}[x_i, y_i] \end{bmatrix} \tag{S18}$$

Here, $x_i$ and $y_i$ describe the location of the $i^{\text{th}}$ pixel of the island under consideration. In the normalized vectors $Col_i$, both the green and blue channel encode the same property (see Equation S7). Therefore, the color vector only contains two independent properties, which encode the relative location in the x direction and the relative location in the y direction of the neuron with respect to other neurons. In the next step, these two properties were extracted:

$$W_i = \begin{bmatrix} \text{Col}_i^{(1)} \\ \frac{1 - \text{Col}_i^{(2)} + \text{Col}_i^{(3)}}{2} \end{bmatrix} \tag{S19}$$

We call the 2D space in which the vectors $W_i$ lay the color space. After placing each $W_i$ in the colorspace, we determined for each element the distance $D_i$ to the $5^{\text{th}}$ closest $W_j$. This distance describes how frequent the color of the pixel is.

$$D_i = \text{sort}\big(\{\sqrt{(x_i - x_j)^2 + (y_i - y_j)^2 \text{ — } \forall (x_j, y_i) \text{ in island}}\}\big)_5 \tag{S20}$$

In Fig. S8 we give a graphical explanation of what the above mentioned distance is defined.

Based on all $D_i$, we chose a threshold $\delta$ such, that one third of them where smaller than $\delta$ and two thirds bigger than $\delta$. We discarded all elements $W_i$ of an island for which $D_i > \delta$. By doing so, we could only keep $W_i$

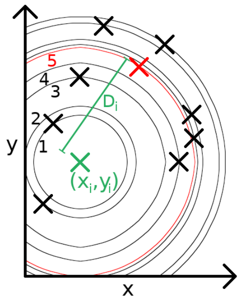


Figure S8: **Distance metric of point in colorspace:** The distance $D_i$ for the $i^{\mathrm{th}}$ point (green) in the colorspace is defined as the euclidean distance to the $5^{\mathrm{th}}$ closest other point (red).

that are similar in the colorspace. While this pre-segmentation of the colorspace is not necessary, it simplifies the subsequent clustering.

Of the remaining color vectors $W_i$, the largest number $n$ was found, for which all $n$ mean locations of an $n$ component gaussian mixture model (gmm) had an euclidean distance of at least 0.25. It was assumed that an island never had more than 5 neurons ($n \leq 5$). The final number of neurons per island was set to $n$. The location of these neurons was chosen as the mean of the pixel location belonging to each of the gmm gaussians.
